# Predictive links between petal color and pigment quantities in natural *Penstemon* hybrids

**DOI:** 10.1101/2023.04.06.535869

**Authors:** Joshua T.E. Stevens, Lucas C. Wheeler, Noah H. Williams, Alice M. Norton, Carolyn A. Wessinger

## Abstract

Flowers have evolved remarkable diversity in petal color, in large part due to pollinator-mediated selection. This diversity arises from specialized metabolic pathways that generate conspicuous pigments. Despite the clear link between flower color and floral pigment production, studies determining predictive relationships between pigmentation and petal color are currently lacking. In this study, we analyze a dataset consisting of hundreds of natural *Penstemon* hybrids that exhibit variation in flower color, including blue, purple, pink, and red. For each individual hybrid, we measured anthocyanin pigment content and petal spectral reflectance. We found that floral pigment quantities are correlated with hue, chroma, and brightness as calculated from petal spectral reflectance data: hue is related to the relative amounts of delphinidin vs. pelargonidin pigmentation, whereas brightness and chroma are correlated with the total anthocyanin pigmentation. We used a partial least squares regression approach to identify predictive relationships between pigment production and petal reflectance. We find that pigment quantity data provide robust predictions of petal reflectance, confirming a pervasive assumption that differences in pigmentation should predictably influence flower color. Moreover, we find that reflectance data enables accurate inferences of pigment quantities, where the full reflectance spectra provide much more accurate inference of pigment quantities than spectral attributes (brightness, chroma, and hue). Our predictive framework provides readily interpretable model coefficients relating spectral attributes of petal reflectance to underlying pigment quantities. These relationships represent key links between genetic changes affecting anthocyanin production and ecological functions of petal coloration.

## Introduction

Angiosperms show spectacular evolutionary diversity in flower color, and a primary mechanism for this diversity is adaptation to attract pollinators to maximize outcross mating success (Fenster et al. 2004; Phillips et al. 2020). A variety of floral pigments, including anthocyanins, betalains, and carotenoids, have evolved through elaboration of specialized metabolic pathways (Grotewold 2006). Pigments confer color by absorbing light from certain wavelengths and scattering or reflecting the remaining light, producing characteristic reflectance spectra defined by the absorbance profiles of pigments. These spectral properties are then perceived and interpreted by pollinators, and selection has favored the evolution of floral pigments that are highly detectable to the visual systems of important pollinators such as bees, birds, butterflies, and moths (e.g., Dyer et al. 2012; Shrestha et al. 2013). While pH, petal thickness, and the presence of metal ions can also be relevant, pigment composition and quantity are usually the key determinants of floral reflectance spectra (Grotewold 2006; Takeda 2006).

The spectral properties of reflected light are often summarized by attributes that include brightness, chroma, and hue. Brightness describes the intensity of the reflected light and is calculated as the total integral of the reflectance spectrum. Chroma describes the saturation of color above greyness for a given brightness, which is determined by the slope of the reflectance curve at its inflection point. Hue provides a description of the peak wavelengths reflected and is typically quantified as an angle in color space (Chittka 1992), which can be calculated using segment classification (Endler and Mielke 2005; Smith 2014). There are straightforward predictions for how pigmentation should affect these spectral attributes. An increase in pigment quantity should reduce brightness and increase chroma, because more pigmentation leads to more light being absorbed (e.g., Papiorek et al. 2013; Van der Kooi 2021). A change in pigment composition (the relative quantities of different pigments) would likely shift floral hue because pigments differ in their absorption spectra.

Anthocyanins, the glycosylated forms of anthocyanidins, are a key class of flora pigments that are widespread across angiosperms and generate a variety of flower colors in flower. Anthocyanidins occur in three different forms that vary in their level of hydroxylation on the B-ring of the molecule: pelargonidin has one hydroxyl group, cyanidin has two, and delphinidin has three. An increase in hydroxylation shifts the absorption spectra of the molecule, making a bluer color: pelargonidin-based anthocyanins tend to appear red, cyanidin-based anthocyanins can show a range of colors from magenta to bluish-purple, and delphinidin-based anthocyanins tend to appear blue (Tanaka et al. 1998).

Evolutionary transitions in flower color are common, resulting in closely related species that differ in substantially in this phenotype. Comparative studies have confirmed that increased pigment amounts result in more saturated flower colors, whereas differences in pigment compositions result in different floral hues (reviewed by Rausher 2008; Sobel and Streisfeld 2013; Wessinger and Rausher 2012). A pervasive assumption is that changes to pigment quantity and composition should affect petal spectral reflectance in a predictable manner. However, to our knowledge, a quantitative framework relating pigmentation and spectral reflectance has not yet been reported. A predictive framework would not only illuminate the underlying mechanisms of petal spectral reflectance, but also has practical value: petal reflectance data is much easier and less expensive to obtain than measurements of pigment quantities. Therefore, studies assessing the predictive relationship between pigment concentrations and flower color data are necessary.

Multivariate statistical approaches are well-suited to identify the links between petal reflectance attributes and pigment quantities. Various multivariate methods are relevant to problems such as this, for example Multivariate Linear Regression (Rencher and Christensen 2012) and Principal Components Regression (Jolliffe 1982), as well as more sophisticated machine learning approaches including Support Vector Regression (Cortes and Vapnik 1995) and Random Forest Regression (Liaw and Wiener 2002). Here we use Partial Least Squares (PLS) regression (Gerlach et al. 1979; Wold et al. 1984; Wold et al. 2001), also known as Projection onto Latent Structures (Abdi 2010), to assess the ability of pigment quantities to predict spectral properties and vice-versa. In PLS, the multivariate dependent and independent datasets are transformed to maximize covariance between them. This approach yields a set of coefficients and rules for transformation that provide an easily interpretable method for predicting a multivariate *Y* matrix from the multivariate *X* input, in our case providing interpretable relationships between pigment quantities and spectral data. An ideal dataset to examine these links is a large one, with substantial variation in floral hue, intensity, and pigment content. One can directly assess the accuracy of model predictions by training the model on a subset of the large dataset and then predicting values for the withheld portion. This process can be done iteratively with random sampling. The coefficients and loadings of the individual variables on the latent structures identify which features contribute most to the prediction.

Here we leverage a phenotypically variable hybrid population of *Penstemon* wildflowers that exhibit substantial variation in flower color. We develop a predictive model relating petal reflectance and anthocyanin pigment quantities, as measured through Thin Layer Chromatography (TLC) methods. We find correlations between spectral attributes (brightness, chroma, and hue) and pigment quantities. Our PLS results suggest this approach is promising for identifying links between pigment quantities and reflectance spectra. Matching well-established assumptions, we find pigment quantities successfully predict spectral attributes, especially hue. Furthermore, analysis of the full reflectance spectra enables accurate inference of pigment quantities.

## Methods

### Study system

The North American angiosperm genus *Penstemon* shows substantial variation in floral pigment production reflecting adaptation to different pollinators. Most *Penstemon* species have blue or purple flowers and are pollinated by bees or wasps (Wilson et al. 2004). Red flowers reflecting adaptation to hummingbird pollination have evolved independently in multiple lineages (Wessinger et al. 2019; Wilson et al. 2007). These flower color differences are due to differential production of anthocyanin pigments: blue- or purple-flowered *Penstemon* species tend to produce anthocyanins based on delphinidin (or a mix of delphinidin and cyanidin), whereas red-flowered *Penstemons* produce pelargonidin-based anthocyanins (Scogin and Freeman 1987; Wessinger and Rausher 2015).

*Penstemon eatonii* (hummingbird-adapted flowers) and *P. laevis* (bee-adapted flowers) are sister taxa (Wessinger et al. 2019) that have diverged in pollination mode and flower color: *P. eatonii* has red flowers due to pelargonidin-based pigments and *P. laevis* has blue flowers due to a combination of delphinidin- and cyanidin-based pigments (Fig. S1). The two species have overlapping distributions in southern Utah and the northern Arizona strip. Where they co-occur, they hybridize due to incomplete reproductive isolation (Crosswhite 1967; Crump et al. 2020). This includes the formation of extensive hybrid zones that exhibit substantial segregating variation in flower color (Crump et al. 2020), varying from the intense red that characterizes *P. eatonii* to the light blue that characterizes *P. laevis*. Hybrids between these species have been given the taxon name *P. x jonesii* (Pennell 1920).

### Field sampling

During May of 2022, we sampled 330 *P. x jonesii* individuals in three geographically proximal populations in canyon systems in Mohave County, Arizona: western Rosy Canyon (n = 161), eastern Rosy Canyon (n = 84), and Potter Canyon (n = 85). We collected fresh flowers by clipping at the pedicel and placing into 25 ml snap-cap tubes that we stored in a cooler backpack with blue ice for later measurement of reflectance spectra (approximately 4-8 hours later). Flowers remain intact for at least 24 hours in this condition. A single flower per individual was used for both reflectance spectroscopy and floral pigment concentrations.

### Petal reflectance measurements

We used reflectance spectroscopy to measure external petal color on the upper lobe surface. We used a Flame T UV-vis spectrometer (Ocean Insight, Dunedin, FL USA) equipped with a Y-shaped reflection probe (Ocean Insight) that supplied light from a deuterium/tungsten light source (Ocean Insight). We calibrated reflectance mesaurements with a spectralon reflectance standard (Ocean Insight). We recorded reflectance data from 300-800 nm using Ocean View software (Ocean Insight) in a darkened room with no incidental light. We sandwiched petal tissue between two cleaned glass microscope slides to eliminate spectral variation due to differences in petal surface curvature. During measurements, the slide was placed directly on top of the upward-oriented light source. This orientation eliminates any backscatter from an underlying surface. Following reflectance measurements, flowers were desiccated on silica gel and transported to University of South Carolina for pigment extraction.

### Calculation of spectral attributes

We processed raw reflectance spectra using the R package *pavo* (Maia et al. 2019). We removed negative values, normalized spectra to range from 0–1, processed spectra using a loess smoothing function with parameter set to 0.2 to reduce noise, and binned spectra into 500 single-wavelength bins from 300–800 nm. We then used the resulting standardized spectra to calculate hue and chroma using a segment classification approach (Smith 2014) and calculated brightness as the total area under the curve from 400–700 nm. Additionally, we calculated hexadecimal color codes of hybrid individuals for plotting using binned spectra and the *pavo* function *spec2rgb*.

### Anthocyanidin pigment extractions and chromatography

We weighed each dried flower on an analytical balance prior to pigment extractions to find flower mass. Next, we extracted anthocyanidins from the flower according to Harborne (1984). Briefly, we submerged petal tissue in 2N HCl overnight, heated the sample to 100°C for 30 minutes to remove sugar moieties, and then performed organic extractions with isoamyl alcohol. We eluted the extracted anthocyanidins in 20 μl of 99% methanol:1% HCl and stored them in -20°C. We used TLC on Cellulose F glass plates (Millipore, Burlington, MA) to separate anthocyanin pigments in Forestal solvent (glacial acetic acid/water/HCl 30:10:3) (Harborne 1967). We spotted 12 μl of each sample for TLC.

To make standard solutions of anthocyanidins, we diluted pelargonidin, cyanidin, and delphinidin standards (Millipore) to a concentration of 1 mg/mL and combined these in equal portions to create a 1:3 multi-anthocyanidin standard solution. We repeated this process to obtain a replicate standard solution. Then we performed a dilution series for each replicate standard with the following levels: 1:6, 1:12, 1:24, 1:48, and 1:96. We ran out two technical replicates of each standard dilution on TLC.

### Pigment quantity calculations

We calculated pigment quantities from TLC plates based on comparisons to our standard dilution series. We photographed TLC plates in a light box using standardized camera settings and lighting conditions (Fig. S2). Specifically, we used a Nikon z50 camera (shutter speed: 1/25, aperture: f16, Iso: 500) equipped with a 550D LED ring light. We imported photos into the program *qTLC* (Mac Fhionnlaoich et al. 2018). Using *qTLC*, we cropped photos, inverted colors, and selected lanes and bands for each plate. Finally, we manually highlighted each pigment spot (Fig. S2). We set the minimum peak prominence (MinPeakProm) parameter to 6 and the band boundaries parameter (divisor) to 1.4 in order to capture the entirety of our pigment spots. This method provided spot intensity values for standards and experimental samples.

We averaged spot intensity values across the technical replicates of each standard dilution series, giving us a single intensity value per dilution level per series. We converted the units from mg/mL (mass concentration) by multiplying each value by the mL loading volume. These replicate dilution series were highly similar (Table S1). We then fit a regression line through these values to find the relationship between spot intensity and anthocyanin pigment quantity (Fig. S3) using the *lme4* R package (Bates et al. 2014). For each sample, we converted spot intensities to pigment quantities by comparison to the standard curve, scaling these values up to the full elution volume (Table S2). We divided the total pigment quantity by the dry flower weight, giving us a total pigment mass fraction expressed in mg/g (mg pigment quantity)/(g dry flower mass). We calculated the fraction of pelargonidin, cyanidin, and delphinidin by dividing each respective pigment mass fractions by the sum of all three.

### Principal components analysis

We performed principal components analysis (PCA) on a dataset that includes the three spectral attributes (brightness, chroma, and hue) and four pigment quantities (fraction pelargonidin, fraction cyanidin, fraction delphinidin, and total pigment mass fraction) for each sampled flower. We used the *prcomp* function from the *stats* package in R to scale and center the data and perform the PCA.

### Partial least squares analysis to relate reflectance spectra to pigment quantities

We conducted three PLS analyses on our data to relate reflectance spectra to pigment quantities: (PLS-A) an analysis to predict spectral attributes from pigment quantities, (PLS-B) an analysis to infer pigment quantities from spectral attributes, and (PLS-C) an analysis to infer pigment quantities from the full floral reflectance spectra. For PLS-A, we used the matrix of pigment quantities (total mass fraction, fraction pelargonidin, fraction cyanidin, and fraction delphinidin) as a multivariate independent data matrix (*X*) and spectral attributes (brightness, chroma, and hue) as the multivariate dependent data matrix (*Y*). For PLS-B, we reversed the direction of PLS-A and used the matrix of spectral attributes as the independent *X* and the matrix of pigment quantities as the dependent *Y*. For PLS-C, we normalized full spectra to the same 0-1 scale, smoothed the data to 1 nm resolution, and treated these data as the multivariate independent data matrix (*X*), where the reflectance intensity value at each nm is treated as a separate individual *x* variable. We treated the matrix of pigment quantities as the dependent *Y*.

For each analysis, we standardized both the *X* and *Y* datasets using the *StandardScaler* in *scikit-learn* (Pedregosa et al. 2011) prior to fitting the PLS model, because initial testing indicated improved performance with transformed data. We optimized the number of PLS components (latent structures) using a 10-fold cross-validation approach. We iterated over a range of component numbers (1-30), calculated the mean-squared error (MSE) of prediction (comparing observed vs. predicted values) for a 10-fold cross-validated model, and plotted the MSE as a function of the number of components (Fig. S4). We chose the optimal number of components based on the inflection point in the MSE curve, finding 2 components was optimal for both PLS-A and PLS-B and 12 components was optimal for PLS-C.

For each mode, we assessed performance using a repeated test/training dataset split. We randomly split the data into halves twenty times, fit the PLS model to the training set, and then used the model to predict values for (1) the samples in the training set and (2) the samples in the test set. We calculated a regression coefficient between the observed true and predicted values for each iteration using *linregress* in *scipy*.*stats* (Virtanen et al. 2020). We report model performance as the average strength of correlation (R^2^) between the observed “true” values vs. the predicted values in the test datasets, averaged over the 20 repeated data splits. We then plotted the distributions of the correlation (R^2^) values for the training and test set predictions, along with the median values. We assessed the contributions of individual *x* variables to the prediction of *Y* by refitting the PLS model to the entire dataset, using the maximum amount of data available, and examining the resulting coefficients that describe the relationship of each *x* variable to each *y* variable.

### Comparison to Random Forest regression

We compared the results of our PLS-C model analysis to another commonly-used machine learning approach: Random Forest (RF) regression. We repeated the steps for PLS-C using RF regression using the *RandomForestRegressor* in *scikit-learn*. We optimized the maximum depth hyperparameter by examining the MSE of prediction as a function of the *max_depth* value. The RF regression did not outperform the PLS-C on test dataset predictions – while predictive abilities were similar, the RF showed signs of overfitting (Fig. S5). In addition, we view the PLS results as more readily interpretable: while the RF fit results do report the relative importance of *x* variables on model predictions, the PLS model returns a set of coefficients that map the relationships between individual *x* and *y* variables, making comparisons of predictive effects within and across models straightforward. Due to these considerations, we chose to use PLS for the primary analysis.

## Results

### Spectral attributes and pigment contents of flowers

Hybrid individuals showed substantial variation in anthocyanidin composition and quantities. All three of the major anthocyanidins (pelargonidin, cyanidin, and delphinidin) were found in the population samples. Methylated anthocyanidins were not detected, consistent with previous studies that have not detected these pigments in *Penstemon* flowers (Scogin and Freeman 1987; Wessinger and Rausher 2015). Individual flowers varied substantially in total pigment quantity, and mostly produced just one or two anthocyanidins that differ by one hydroxyl level (i.e., pelargonidin with cyanidin or cyanidin with delphinidin; Fig. S6). Accordingly, we observed substantial variation in spectral attributes of reflectance data (Fig. S6).

We used a PCA to visualize the relationship between pigment quantities and spectral attributes (Fig. 1). The first two principal components explained 77.6% of the variation within the dataset. PC1 largely separates samples with higher hue values (bluer colors), greater delphinidin fractions, and smaller pelargonidin fractions from those with lower hue values (redder colors), smaller delphinidin fractions, and greater pelargonidin fractions. PC2 separates samples with greater total pigment mass fraction, greater cyanidin fractions, and lower brightness from samples with small total pigment fraction and higher brightness.

**Figure 1.**
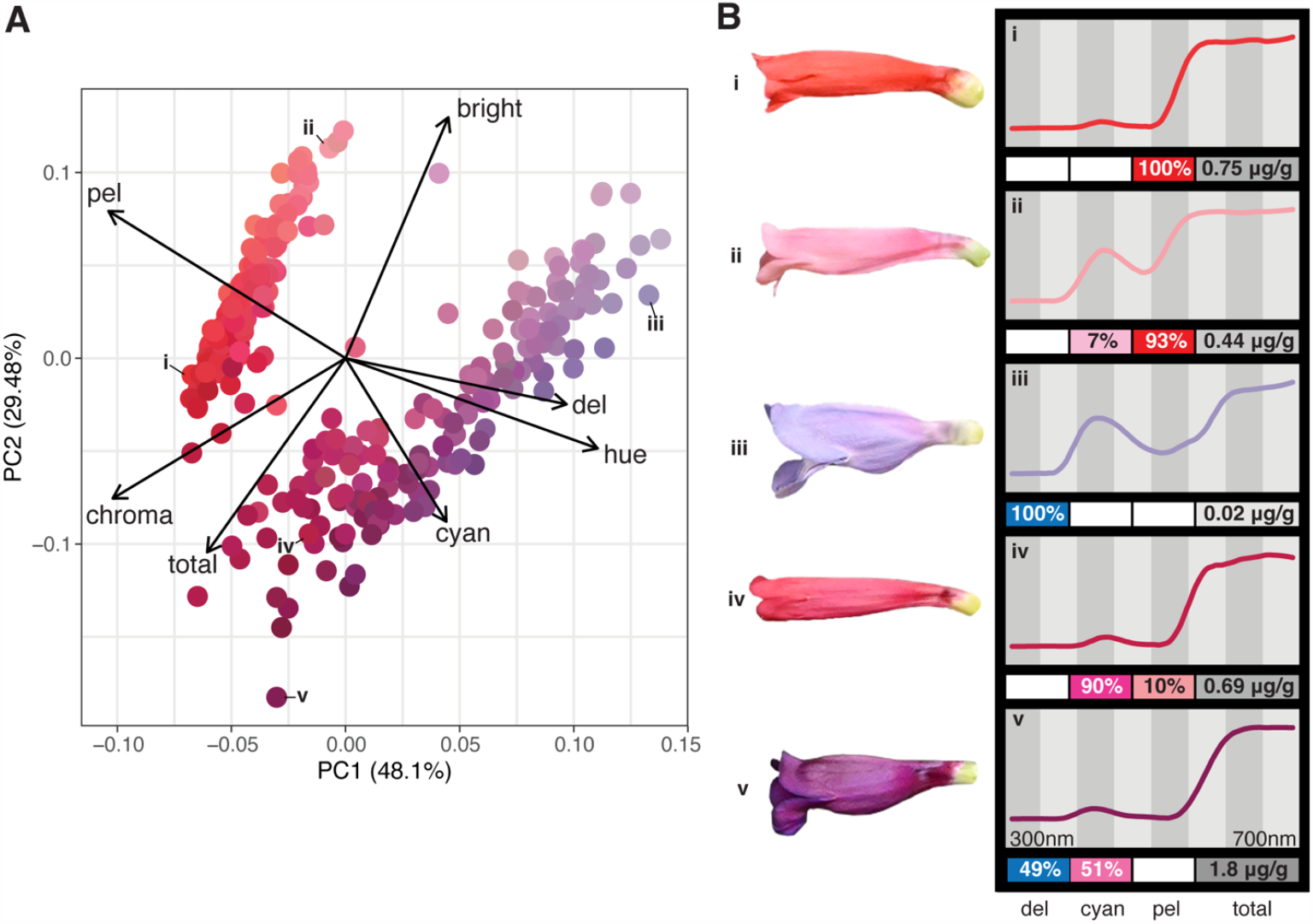
Relationships between spectral attributes and pigment quantities. A: PCA plot of *P. x jonesii* flowers based on their spectral attributes and pigment quantities. Arrows represent loadings and colors show hexadecimal color codes based on individual spectra. B: Photographs, reflectance spectra, and pigment quantities for representative individuals. Bright: brightness, cyan: fraction cyanidin, del: fraction delphinidin, total: total pigment mass fraction, pel: fraction pelargonidin, μg/g: micrograms pigment per gram dry flower weight.

### PLS is a promising approach to relate pigment quantities to reflectance spectra data

We fit three PLS models designed to predict spectral attributes from pigment quantities (PLS-A), infer pigment quantities from spectral attributes (PLS-B), and infer pigment quantities from the complete reflectance spectra (PLS-C). For all models, we compared model performance on training and test datasets and found that the test set performance approached the training set performance for all variables, although the test set performance was usually slightly lower than the training set for PLS-C (Fig. 2, Fig. 3). When fitting models to the entire dataset, the R^2^ between predicted and observed values increases, as expected (Table S3). The similar performance of the test vs. training sets implies a limited effect of overfitting the model to the training data. Overall these results suggest PLS is a promising approach to generate informative links between spectral reflectance and pigment datasets.

**Figure 2.**
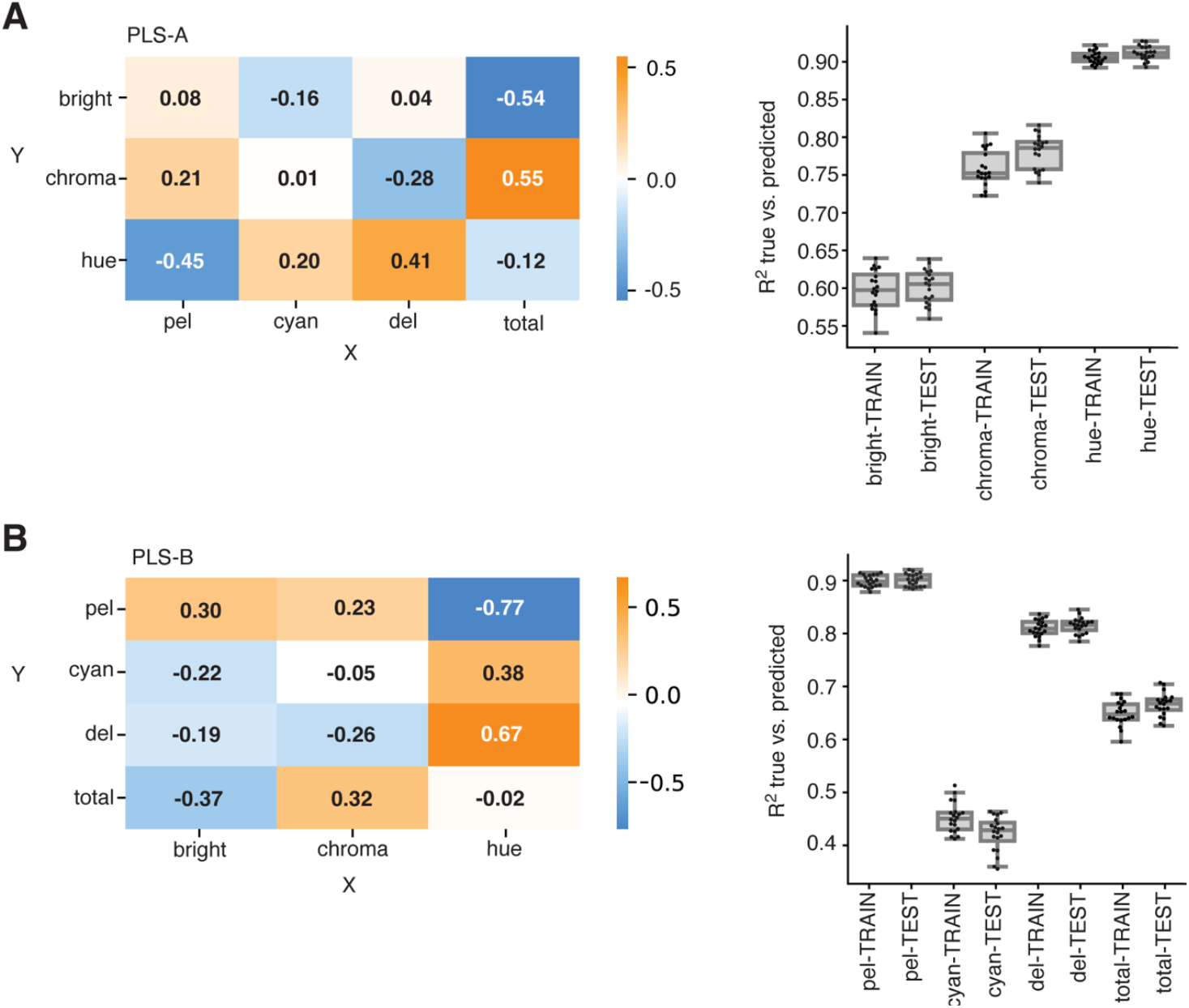
Results of PLS-A and PLS-B models relating spectral attributes and pigment quantities. A: PLS-A, B: PLS-B. Heatmap at left shows the model coefficients relating *x* and *y* variables. Boxplots at right show the distributions of correlation coefficients of true vs. predicted values averaged across 20 replicate data splits. Bright: brightness, cyan: fraction cyanidin, del: fraction delphinidin, total: total pigment mass fraction, pel: fraction pelargonidin.

**Figure 3.**
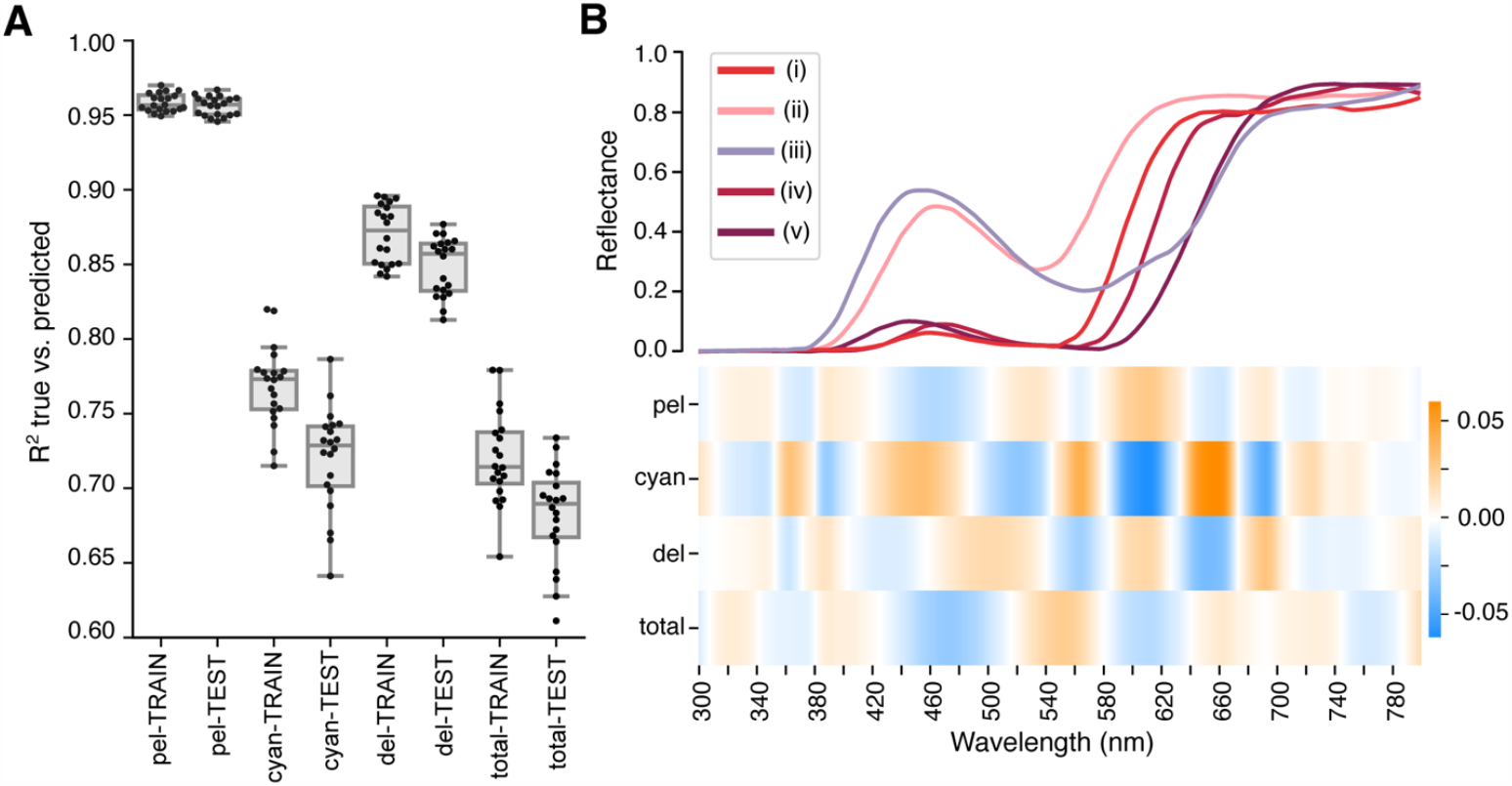
Results of PLS-C model relating spectral attributes and pigment quantities. A: Distributions of correlation coefficients of true vs. predicted values averaged across 20 replicate data splits. B: Top shows spectra for representative individuals shown in Fig. 1B, bottom shows heatmap with the magnitudes of model coefficients relating *x* wavelengths and *y* variables as estimated from the entire dataset. Bright: brightness, cyan: fraction cyanidin, del: fraction delphinidin, total: total pigment mass fraction, pel: fraction pelargonidin.

### Spectral attributes are predicted by pigment quantities

Our PLS-A model predicted spectral attributes from pigment quantities with relative success (Fig. 2A). The average correlation coefficients between predicted and observed spectral attributes in the test datasets were relatively high (Fig. 2A; Table S3), with best performance observed for predictions of hue (mean R^2^= 0.91), followed by chroma (mean R^2^= 0.78), and then brightness (mean R^2^= 0.60). The major contributors to hue predictions were delphinidin fraction (positive effects) and pelargonidin fraction (negative effects), although cyanidin fraction also provided a minor positive contribution. Chroma predictions were predominantly based on total pigment mass fraction (positive effect), as well as pelargonidin (positive effect) and delphinidin (negative effect) fractions. Brightness predictions were largely contributed by total pigment and cyanidin fractions (negative effects).

### Limited inference of pigment quantities from spectral attributes

Our PLS-B model had mixed success inferring pigment quantities from spectral attributes (Fig. 2B). Correlations between predicted and observed pigment fractions in the test datasets were relatively high for pelargonidin (R^2^= 0.90) and delphinidin (R^2^= 0.82) fractions, but lower for the total pigment mass fraction (R^2^= 0.67) and poor for the cyanidin fraction (R^2^= 0.42). The inferred values of delphinidin fractions and pelargonidin fractions were both strongly based on hue: hue contributed positively to predicted delphinidin fractions and negatively to predicted pelargonidin fractions. Hue also had a minor positive contribution to predicted cyanidin fractions. Chroma and brightness contributed negative effects to the predicted delphinidin and cyanidin fractions and positive effects to predicted pelargonidin fraction. Finally, the predictions for total pigment mass fraction were predominantly contributed by chroma (positive effect) and brightness (negative effect).

### Strong signals in full spectral data allows inference of pigment quantities

The PLS-C model based on the full reflectance spectra revealed that the spectra contain a large amount of information that enables relatively accurate inference of pigment quantities. This model performed very well at inferring pelargonidin fractions (mean R^2^= 0.96), followed by delphinidin fractions (mean R^2^= 0.85), cyanidin fraction (mean R^2^=0.72), and total pigment mass fraction (mean R^2^= 0.68). The coefficients of the PLS-C model relate each individual wavelength intensity value to each pigment quantity. These coefficients exhibit a banding pattern resembling a barcode when arrayed along the spectrum (Fig. 3). Each *Y* pigment quantity has a characteristic pattern of positive and negative bands of coefficients. Strikingly, there are several hotspot regions with clear signals associated with relative pigment fractions and total pigment mass fraction. For example, the wavelength band between 580-640 nm show distinct effects (positive vs. negative) on the predicted values of each of the inferred pigment variables (Fig. 3). Similarly, the region between 420-540 nm also shows contrasting effects on each of the pigment quantities. In combination, these distinctive banding patterns yield accurate predictions of pigment composition and quantities.

## Discussion

### PLS models reveal close links between flower color and pigments

The goal of our study was to answer two questions. First, can we predict the spectral attributes of flowers if we know which pigments they produce? Second, can we infer what pigments are being produced by flowers if we know their reflectance spectra? The ability to predict one type of data from the other has practical relevance for empiricists working on flower color. The predictive relationships between pigment quantities and reflectance spectra describe a key link between genetic variation and ecological function. Variation in pigmentation results from differences in the relative expression and activity of pigment producing enzymes, while reflectance spectra are relevant to perception by pollinators.

In our dataset, spectral attributes of *Penstemon* flowers were well-predicted by pigment quantities. In other words, knowing the amounts and relative proportions of anthocyanidins provides a reasonable prediction of spectral attributes. Hue was the best predicted attribute based on pigment data, largely depending on the relative delphinidin vs. pelargonidin fraction of flowers. The strong dependence of hue on relative pigment composition is consistent with the biochemical differences of anthocyanidins – that hydroxylation levels shift the absorption spectra from redder to bluer colors (Tanaka et al. 1998). The cyanidin fraction of flowers had more minor contributions to hue because in our dataset, cyanidin fraction provided little information on hue: cyanidin increases hue relative to pelargonidin but decreases hue relative to delphinidin. We imagine that cyanidin fraction might strongly predict hue in other datasets involving variation in only two pigments, for example, in populations segregating combinations of delphinidin and cyanidin only.

Chroma and brightness largely depended on total pigment content, but were less successfully predicted by pigment content, relative to hue. This may indicate that other petal features segregating in the population affect these attributes, for example the presence of additional pigments or petal texture elements. Our reduced ability to predict brightness and chroma from pigment extracts might arise for technical reasons. First, we suspect there is greater error in our estimates of total pigment content compared to the relative pigment fractions, since evaporation or slight pipetting errors would affect the total amount of pigment (but not relative fractions) applied to TLC plates. Second, our petal reflectance measurements were taken on a single position on the flower, whereas the pigment extractions are based on the entire flower. Any variation in pigment deposition across the flower will weaken the relationship between measured brightness and chroma vs. total pigment mass fraction.

In our dataset, we were surprisingly successful at inferring not only pigment identity, but relative pigment fractions, based on reflectance data. The full spectrum contained much more information than the simpler spectral attributes (brightness, chroma, and hue), including barcode-like wavelength bands that diagnose pigment composition and quantity. Yet, even with the full spectra for flowers, inferences of cyanidin fraction and total pigment content were less accurate, suggesting there is less unique information on these values within the reflectance spectra. Because we normalize raw reflectance spectra, we rely on the shape of the spectra for our predictive models, and there might simply be more information about delphinidin and pelargonidin fractions in the shape of the spectra than cyanidin and total pigment mass fraction. Moreover, the weaker prediction for total pigment content could again reflect greater technical errors in estimates of total pigment mass fraction.

### Promise and limitations of our approach

The PLS modeling approach performed well compared to a random forest regression approach and, importantly, provides easily interpretable coefficients that describe contributions of individual *x* variables to *y* variable predictions. Based on these coefficients, we can find the linear combination of variable effects while accounting for the state of other variables in the dataset. This feature makes PLS ideally suited for disentangling the relationships among spectral and pigment datasets. The degree to which this approach will prove successful in other systems may depend on the biological details and evolutionary context. In our case, individuals in the dataset have a close evolutionary relationship, being hybrids derived from sister species. Individuals vary in anthocyanin content but perhaps have not diverged in other traits that may affect flower color such as differences in co-pigmentation or cellular pH, and additional anthocyanin modifications such as methylation of B-ring hydroxyls (Fournier-Level et al. 2011; Tanaka et al. 2008). Over longer evolutionary timescales, pigment content would not necessarily be inferred from petal reflectance spectra. For example, convergent evolution of flowers that appear red involves multiple different pigment profiles, including anthocyanin and carotenoid pigment combinations (Ng and Smith 2016). A dataset including such variation may have low success inferring pigment content from reflectance spectra, depending on the number of representative individuals present in the training data.

### Implications for the genetics of flower color differences between *P. laevis* and *P. eatonii*

The pattern of segregating pigment variation in *P. x jonesii* hybrid swarms provides insight into the genetic basis of flower color in this system, to be verified in future molecular genetic studies. The anthocyanin biosynthesis pathway is well-characterized and conserved in angiosperms (Fig. 4). The production of cyanidin and delphinidin requires the activity of the enzymes Flavonoid 3’-hydroxylase (F3’H) and Flavonoid 3’,5’-hydroxylase (F3’5’H), respectively, which add hydroxyl groups onto the B-ring (Holton et al. 1993). Variation in the relative amounts of anthocyanidins can result from altered expression or function of F3’H or F3’5’H or by substrate specialization of the common downstream enzyme dihydroflavonol 4-reductase (DFR) on specific precursors (e.g., specialization on DHK vs. DHM) (Smith and Rausher 2011). Variation in the total pigment amount often results from variation in transcription factors that regulate the structural genes of the pathway, including members of *MYB, bHLH*, or *WDR* families (reviewed by Lloyd et al. 2017).

**Figure 4.**
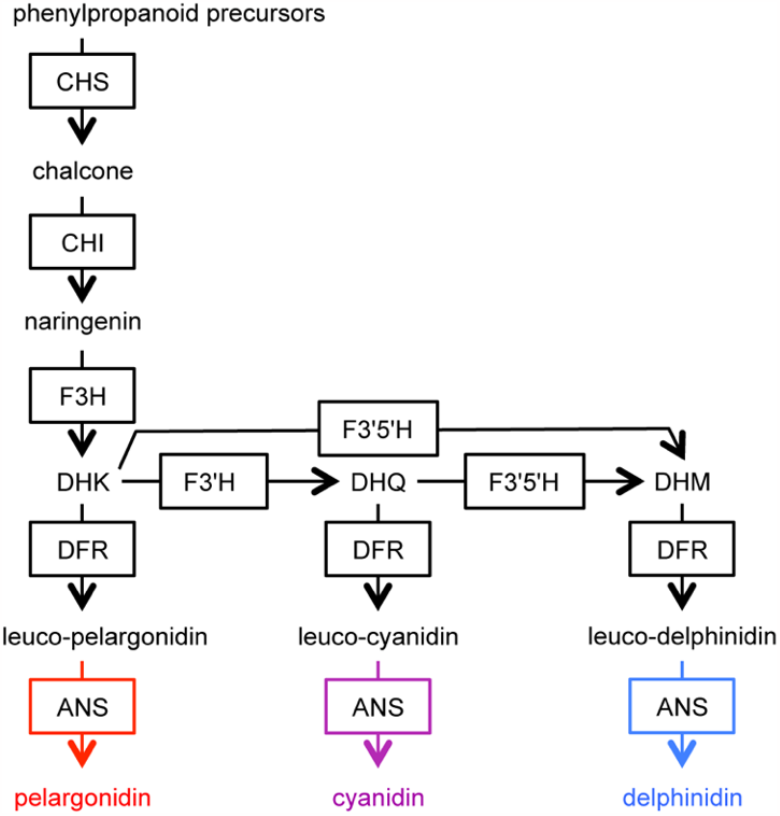
Simplified anthocyanin biosynthesis pathway. Boxes indicate enzymes and arrows indicate enzymatic reactions. ANS: anthocyanidin synthase, CHI: chalcone isomerase, CHS: chalcone synthase, DFR: dihydroflavonol-4-reductase, DHK: dihydrokaempferol, DHM: dihydromyricetin, DHQ: dihydroquercetin, F3H: flavanone-3-hydroxylase, F3’H: flavonoid 3’-hydroxylase, F3’5’H: flavonoid 3’,5’- hydroxylase.

The parent species of *P. x jonesii* differ in both anthocyanidin type and quantity: the blue-flowered *P. laevis* produces delphinidin and cyanidin in modest quantities, whereas the red-flowered *P. eatonii* produces large amounts of pelargonidin. This difference in pigment production is similar to that observed in a previously studied *Penstemon* sister species pair: *P. neomexicanus* has blue-purple flowers due to delphinidin production and *P. barbatus* has red flowers due to pelargonidin production (Wessinger and Rausher 2014). Interestingly, in a genetic cross between *P. neomexicanus* and *P. barbatus*, F2 progeny exhibited just two categories of relative pigment production, although total pigment mass fraction was not quantified. Specifically, individuals produced either delphinidin or pelargonidin, due to allelic differences at the hydroxylating enzyme F3’5’H, with a dominant functional allele associated with delphinidin production and a recessive non-functional allele associated with pelargonidin production (Wessinger and Rausher 2014).

The natural *P. x jonesii* hybrids derived from *P. laevis* and *P. eatonii* show much more variation in relative anthocyanidin production, suggesting the control of pigmentation is more complex than observed in the *P. barbatus–P. neomexicanus* system. Flower color in *P. x jonesii* may involve allelic differences between species in both hydroxylases F3’H and F3’5’H, explaining why each of the three types of anthocyanidins can be found in hybrids. Interestingly, we only observe combinations of anthocyanidins with neighboring hydroxylation levels, which likely reflects competition among enzymes for common substrates. For example, the combination of cyanidin and delphinidin pigments reflects competition for DHQ between DFR and F3’5’H, with the relative proportions of pigments determined by relative expression and function of these two enzymes. Similarly, the combination of pelargonidin and cyanidin reflects competition between DFR and F3’H for the common substrate DHK. Pelargonidin in combination with delphinidin (without cyanidin) is not observed. Perhaps delphinidin production in this system relies on DHQ as a precursor via hydroxylation by F3’H, followed by the addition of the third hydroxyl by F3’5’H, rather than a direct conversion of DHK to DHM by F3’5’H. In this scenario, production of delphinidin and pelargonidin, but not cyanidin, would likely require a delicate balance of enzyme concentrations and may not be possible without additional mechanisms for reducing the production of cyanidin from DHQ. Specific mutations altering the substrate specificity of DFR can also underlie evolutionary shifts in flower color (Smith and Rausher 2011), and could contribute to fine-tuning pigment profiles. However, these mutations likely have a smaller mutational target size and hence arise less frequently than evolutionary changes in the function or expression of hydroxylating enzymes. Such mutations were not found to underlie the flower color difference in the *P. barbatus–P. neomexicanus* system, but a role for DFR substrate specificity is possible in theory.

In addition, at least one additional locus affects total pigment mass fraction. A common type of mutation affecting overall pigmentation is change in function or expression of a transcription factor that promotes or represses pathway enzymes (e.g., Quattrocchio et al. 1999; Schwinn et al. 2006; Streisfeld and Rausher 2011). In addition, the anthocyanin pathway is embedded within the flavonoid pathway and other enzymes compete with hydroxylating enzymes and DFR for common substrates. This includes Flavonol synthase (FLS), which converts DHK, DHQ, and DHM into the flavonols kaempferol, quercetin, and myricetin, respectively (reviewed by Winkel-Shirley 2001). Thus, a mutation affecting the expression or function of FLS can affect levels of anthocyanin pigmentation (e.g., Lüthi et al. 2022). Future genetic studies in the *P. laevis–P. eatonii* system will test these hypotheses.

## Conclusions

Flower color variation within hybrid populations provides an opportunity to examine the links between phenotypic variation in floral spectral reflectance and variation at the underlying biochemical level. The PLS approach used here enables quantification of these links in a predictive framework. Characterizing these links is key to understanding how genetic changes to enzymes in biochemical pathways generate ecologically-relevant phenotypic differences. The observed flower color variation in *P. x jonesii* hybrid zones may have important effects on pollinator visits by bees and hummingbirds. Bees and birds have very different visual systems that differ in spectral sensitivities and thus differ in abilities to distinguish between colors. The flower color differences between the parent species of *P. x jonesii* presumably reflect adaptation to these distinct visual systems. Future work in the *P. x jonesii* system will apply pollinator visual models to determine the functional significance of hybrid flower color variation on pollinator perception, with implications for mating patterns within the hybrid zone and gene flow between parent species.

## Data Availability

Upon acceptance, all data and scripts will be deposited in publicly accessible databases.

## Acknowledgements

We thank Mikel Stevens for support with this project, Benjamin Stone for fieldwork assistance, and the Bureau of Land Management for sampling permissions. Daniel Speiser and Benjamin Stone provided helpful feedback on the manuscript. This work was supported by NSF award DEB 2052904 to C.A.W.

## Notes

### Competing Interest Statement

The authors have declared no competing interest.

