## Supplemental Text for "Predictive links between petal color and pigment quantities in natural *Penstemon* hybrids"

Table S1. Correlations ($r^2$) between anthocyanidin standard dilution series values.

| Pigment | $r^2$ |
|---|---|
| Pelargonidin | 0.9924 |
| Cyanidin | 0.9836 |
| Delphinidin | 0.9723 |

Table S2. Linear models estimated from the standard curve for relating TLC spot intensities to pigment mass, where $pigment\ mass \sim slope \cdot intensity + intercept$

| Pigment | Slope | Intercept |
|---|---|---|
| pelargonidin | 4.648e-09 | -1.686e-05 |
| cyanidin | 4.706e-09 | -9.207e-05 |
| delphinidin | 5.692e-09 | -9.7173-05 |

Table S3. Correlation coefficients ($R^2$) between predicted and true values of *y* variables in PLS models across 20 test and training splits and using the model fit with the entire dataset.

| Model | *y* variable | Mean $R^2$ (training) | Mean $R^2$ (test) | $R^2$ (entire dataset) |
|---|---|---|---|---|
| PLS-A | Brightness | 0.597 | 0.601 | 0.597 |
| PLS-A | Chroma | 0.758 | 0.781 | 0.770 |
| PLS-A | Hue | 0.906 | 0.912 | 0.908 |
| PLS-B | Fraction pelargonidin | 0.899 | 0.901 | 0.900 |
| PLS-B | Fraction cyanidin | 0.452 | 0.422 | 0.440 |
| PLS-B | Fraction delphinidin | 0.810 | 0.815 | 0.812 |
| PLS-B | Total pigment mass fraction | 0.649 | 0.666 | 0.657 |
| PLS-C | Fraction pelargonidin | 0.959 | 0.956 | 0.960 |
| PLS-C | Fraction cyanidin | 0.769 | 0.720 | 0.762 |
| PLS-C | Fraction delphinidin | 0.870 | 0.849 | 0.867 |
| PLS-C | Total pigment mass fraction | 0.720 | 0.682 | 0.715 |

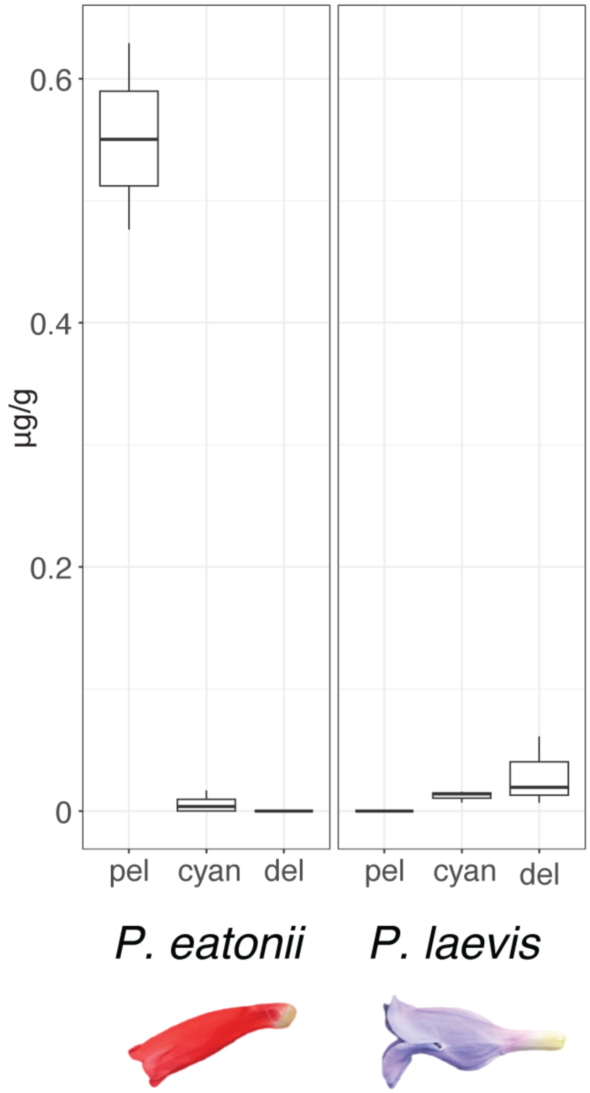


Figure S1. Anthocyanidin content of *P. laevis* and *P. eatonii* flowers. Shown are pigment mass fractions expressed in micrograms (μg) anthocyanidin / grams (g) dry weight. Cyan: cyanidin, del: delphinidin, pel: pelargonidin.

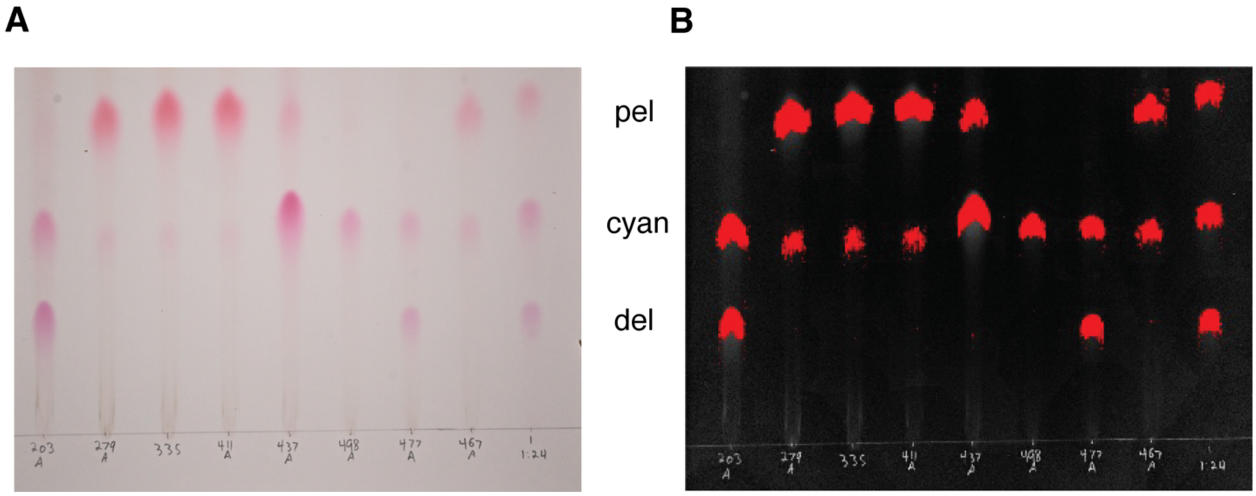


Figure S2. Approach for quantifying pigment quantities from thin layer chromatography (TLC) plates. A: Photograph of representative sample TLC plate. B: Analysis of spot intensity using *qTLC*. Cyan: cyanidin, del: delphinidin, pel: pelargonidin.

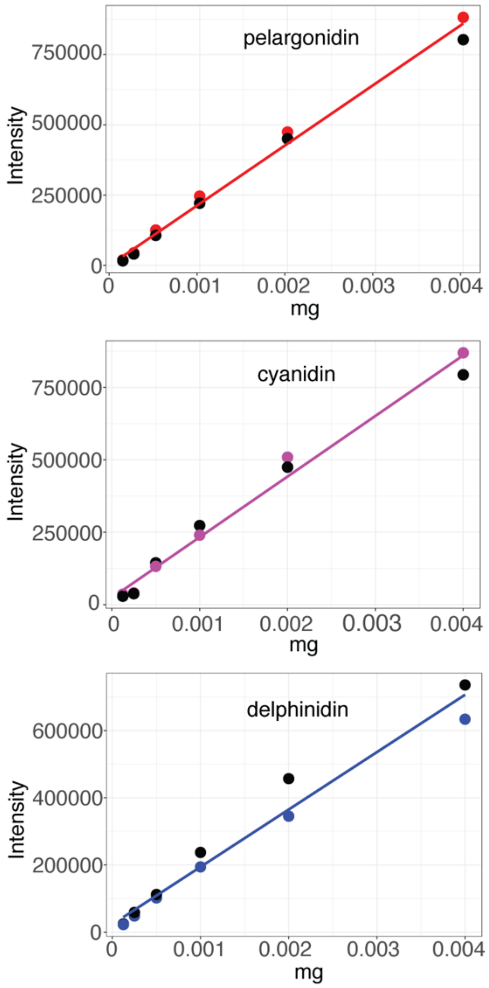


Figure S3. Relationship between TLC spot intensity and anthocyanidin quantities in milligrams (mg) estimated from standard dilution series.

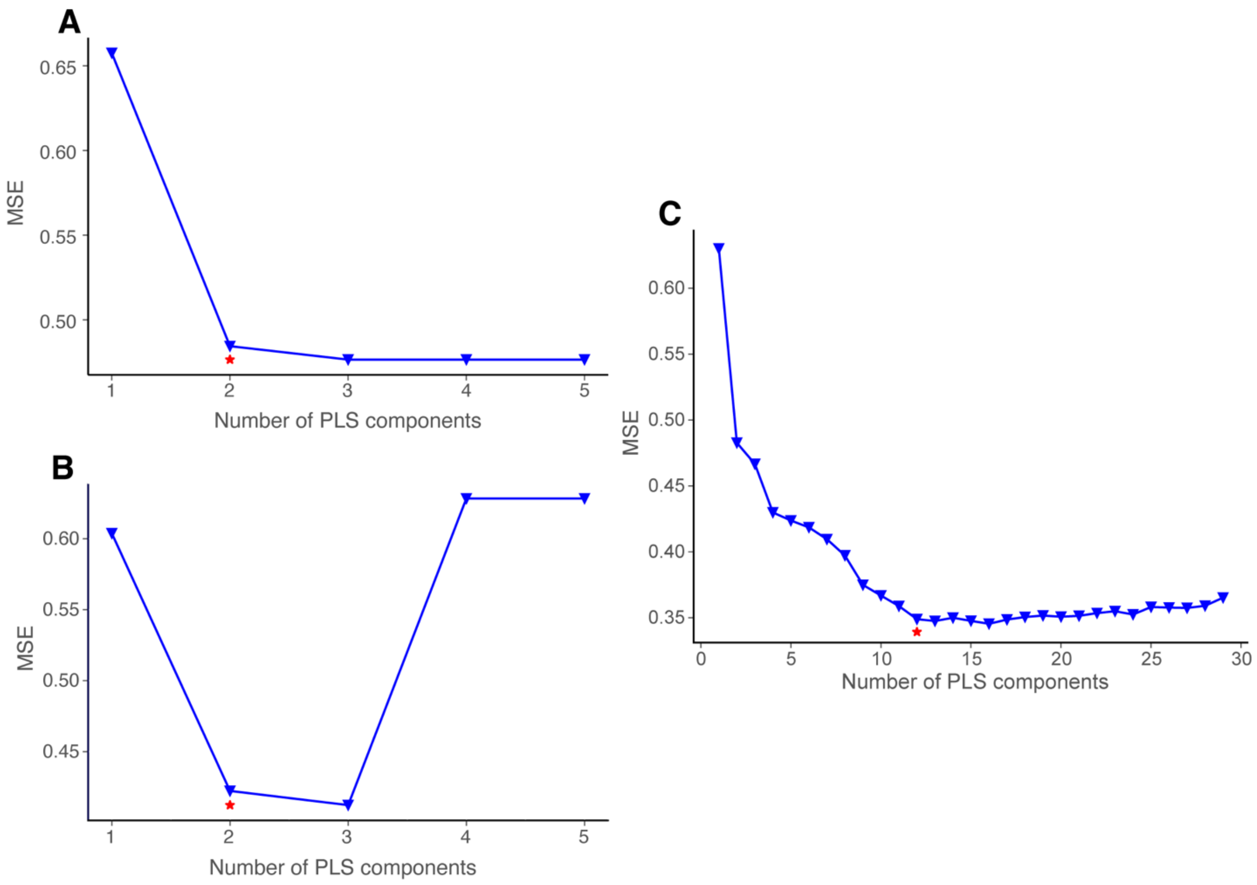


Figure S4. The mean-squared error (MSE) of prediction (comparing true vs. predicted values) for 10-fold cross-validated models as a function of the number of components. Stars indicate optimal number of components based on the inflection point in the MSE curve, where the MSE first reaches its appropriate minimum value.

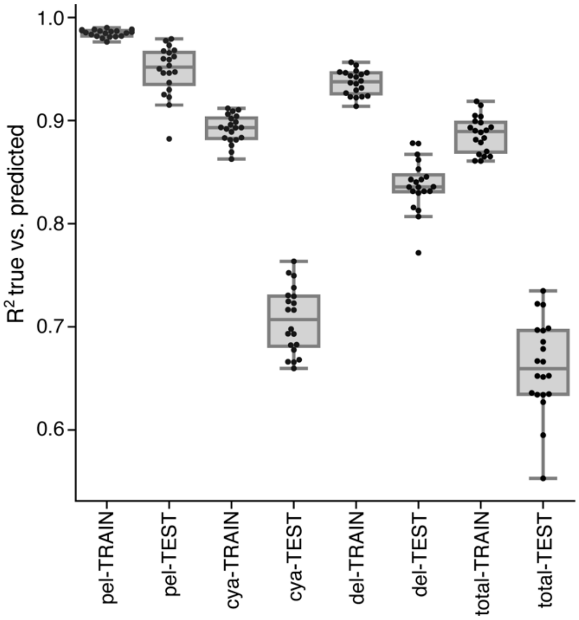


Figure S5. Comparison of random forest (RF) regression to the PLS-C model. Shown are the mean-squared error (MSE) of prediction (comparing true vs. predicted values) for the RF model with the same 20 test-training splits used to assess PLS-C. Cya: cyanidin, del: delphinidin, pel: pelargonidin.

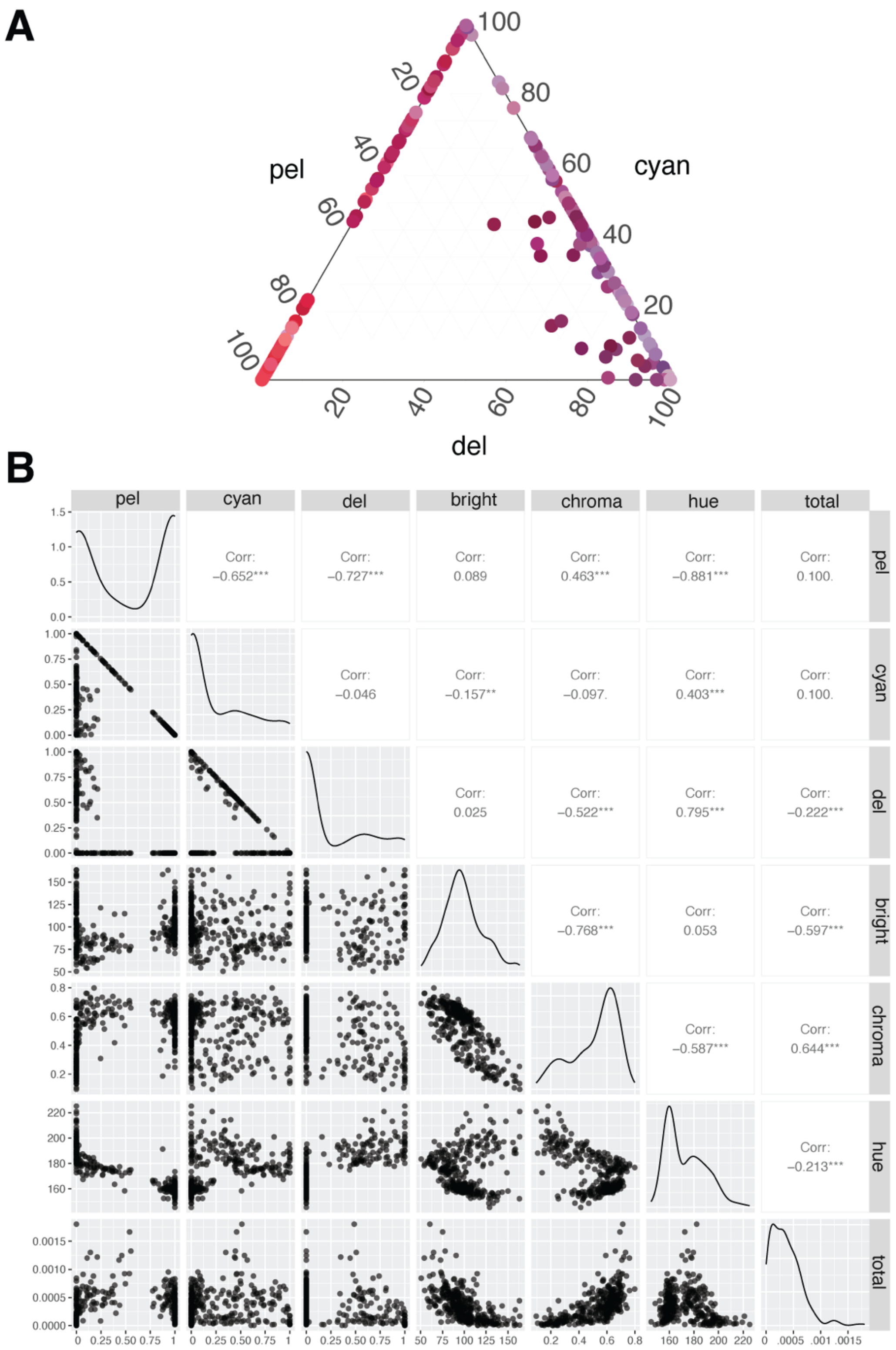


Figure S6. Variation in pigment quantities and spectral attributes across samples. A: triangle plot showing relative fractions of anthocyanidins, B: pairwise scatterplots showing relationships between the three spectral attributes and four pigment fractions. Bright: brightness, cyan: cyanidin, del: delphinidin, pel: pelargonidin.
